## Supplemental Materials for "An Optimized CRISPR/Cas9 Approach for Precise Genome Editing in Neurons"

### Figure supplements

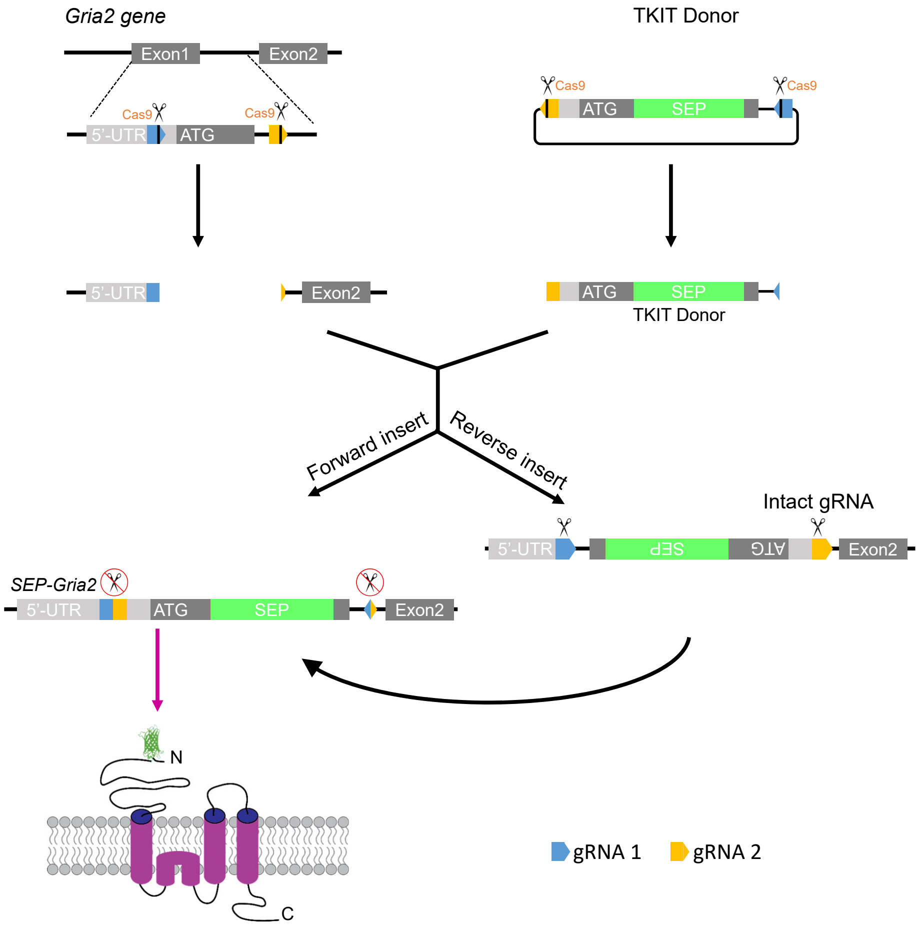

**Figure supplement 1. Strategy to promote donor DNA insertion in the correct orientation**

Graphical representation of TKIT-mediated genome editing with *Streptococcus pyogenes* Cas9. The blue and orange pentagons indicate guide RNA 1 and guide RNA 2 recognition sites, respectively. The black line within the pentagon indicates the Cas9 cleavage site. The light gray rectangle indicates the 5-UTR of gene of interest and the dark gray rectangle the coding sequence. Black arrows indicate the possible outcomes following genomic insertion of the donor DNA.

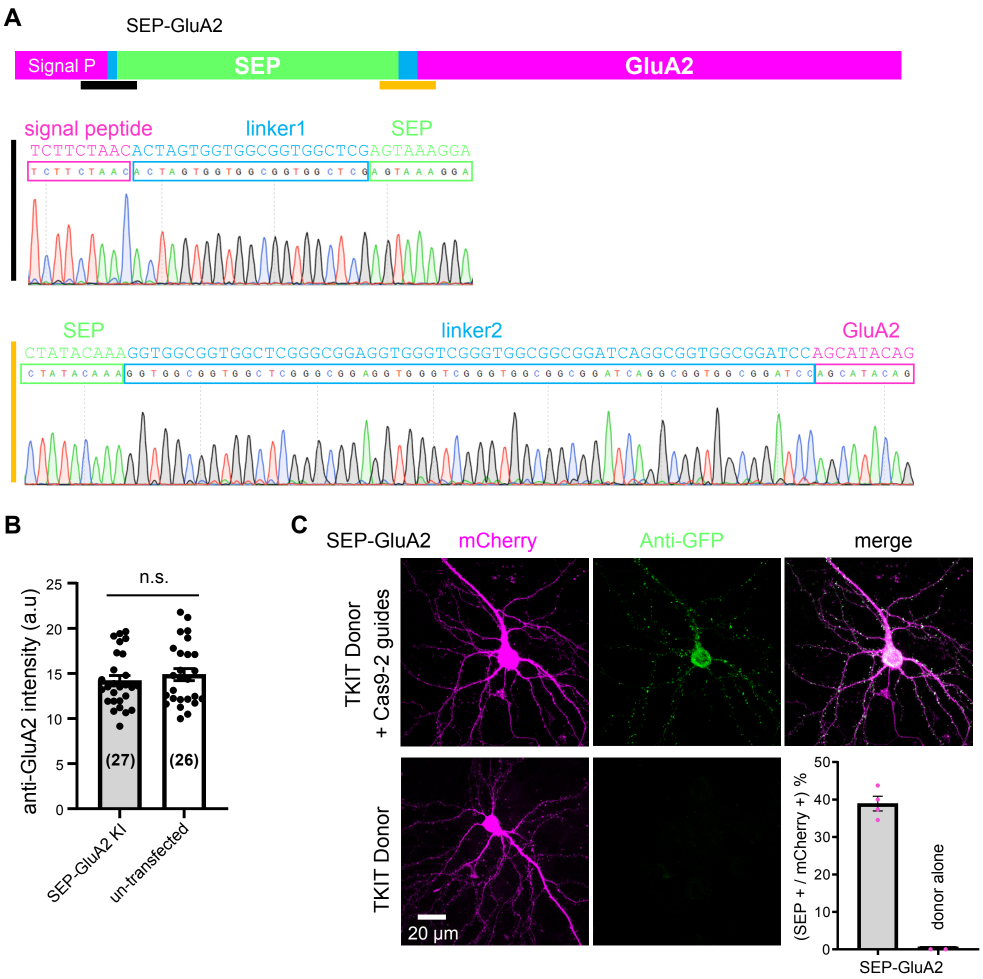

**Figure supplement 2. Correct splicing of SEP-GluA2 and comparable GluA2 levels in KI and WT surrounding neurons**

(A) Sequence analysis of cDNA reverse-transcribed from mRNA extracted from neuronal cultures electroporated with TKIT SEP-GluA2 constructs. The black line indicates the junction between the signal peptide and linker1-SEP and the orange line indicates the junction between SEP-linker2 and the mature protein. The magenta region indicates the endogenous protein sequence, the green sequence indicates SEP and the blue sequence indicate linkers. (B) Quantification of synaptic GluA2 levels with immunofluorescence. GluA2 levels in SEP-GluA2 KI neurons are comparable with non-transfected (WT) neurons (data from 27 SEP-GluA2 KI neurons and 26 surrounding non-transfected neurons). Data presented as means ± SEM. Individual data points are shown as black dots. Unpaired Student’s t-test for comparison ($p=0.4495$). (C) Transfection of TKIT donor alone does not produce KI. Representative images showing TKIT of SEP-GluA2 KI compared to TKIT donor-only control. Average efficiency of KI for SEP-GluA2 compared to donor-only control (bottom right corner). Scale bar indicates 20 µm.

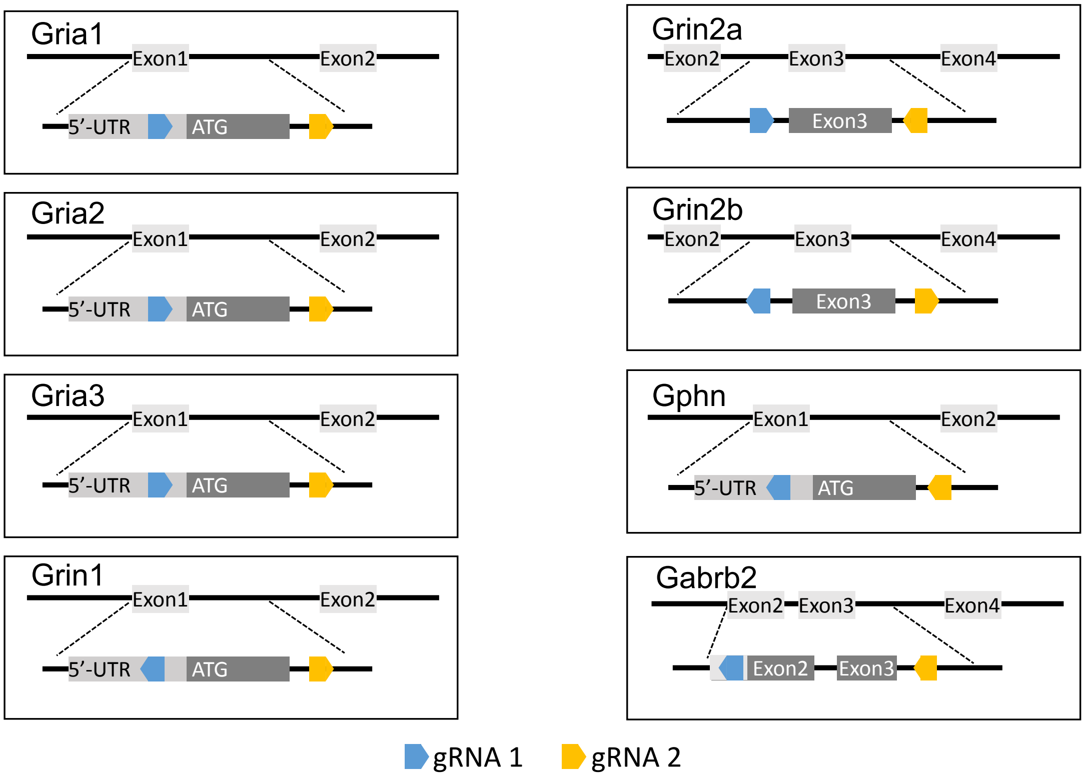

**Figure supplement 3. Location of TKIT guide pairs**

Schematic illustrating the location of all guide RNAs and their orientation in all genes targeted for TKIT. In each panel the top portion indicates the partial exons and introns being targeted. The bottom part shows an enlargement of the region to be edited. The light gray rectangle indicates the 5’-UTR. The dark gray rectangle indicates the coding sequence. The blue and orange pentagons indicate guide RNA (gRNA) 1 and 2, respectively. The genes *Gria1*, *Gria2*, *Gria3*, *Grin1*, *Grin2a*, *Grin2b*, *Gepn* and *Gabrb2* encode for GluA1, GluA2, GluA3, NR1, NR2A, NR2B, Gephyrin and $GABA_AR$ Beta-2, respectively.

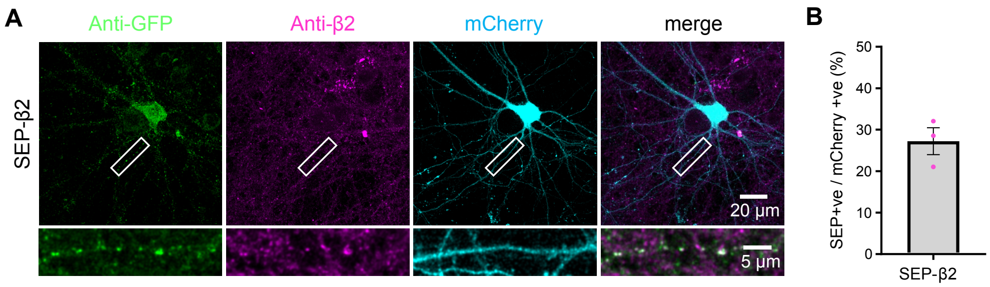

**Figure supplement 4. TKIT of SEP-GABA$_A$R β2**

(A) Representative immunofluorescent images demonstrating TKIT of SEP-GABA$_A$R β2 (SEP-β2). The dendritic region within the white box is enlarged below to better illustrate co-localization between SEP-β2 (green) and anti-β2 signal (magenta). Note: the cell-fill channel was removed from the enlarged overlay images for clarity. Scale bars display 20 μm or 5 μm. (B) Average efficiency of SEP-β2 KI was 27.22±3.24%. Data presented as mean ± SEM.

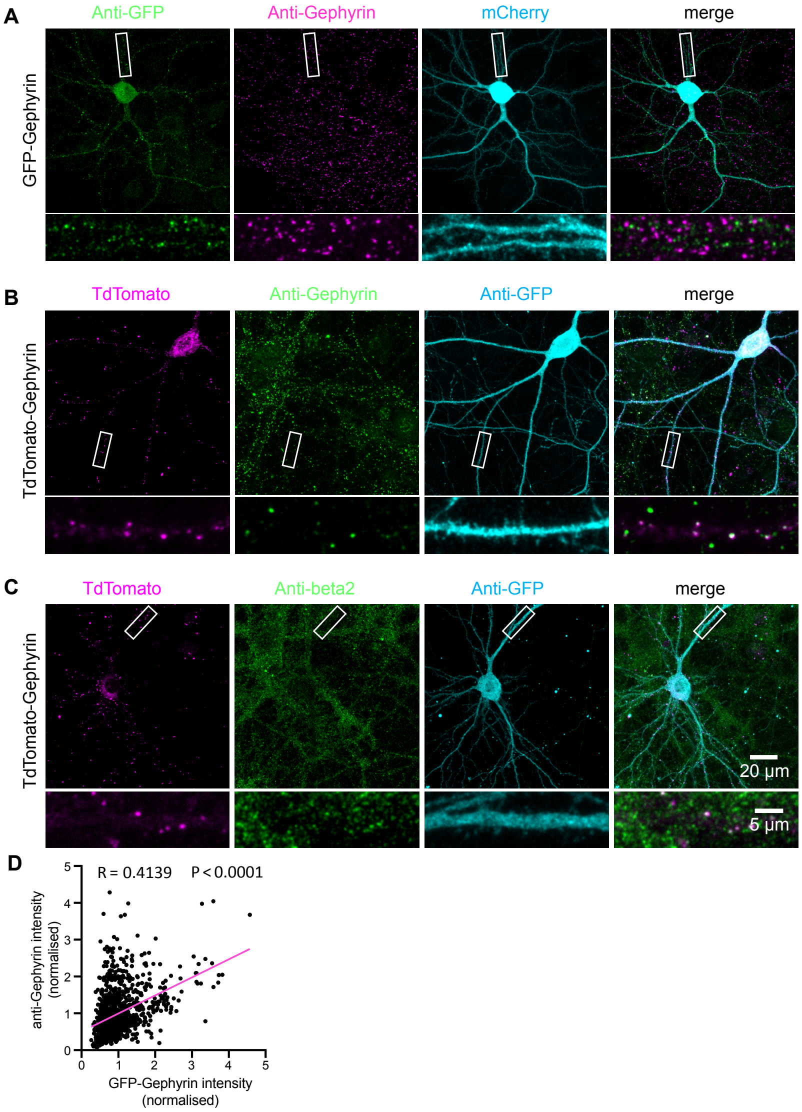

**Figure supplement 5. TKIT of GFP-Gephyrin and tdTomato-Gephyrin**

(A) Representative immunofluorescent images demonstrating TKIT of GFP-Gephyrin. The dendritic region within the white box is enlarged below to better illustrate co-localization between GFP-Gephyrin (green) and anti-Gephyrin signal (magenta). (B) Representative immunofluorescent images demonstrating TKIT of tdTomato-Gephyrin. The dendritic region within the white box is enlarged below to better illustrate co-localization between tdTomato-Gephyrin (magenta) and anti-Gephyrin signal (green). (C) Representative immunofluorescent images demonstrating TKIT of tdTomato-Gephyrin. The dendritic region within the white box is enlarged below to better illustrate co-localization between tdTomato-Gephyrin (magenta) and the β2 $GABA_A$ receptor subunit (green). Note: the cell-fill channel was removed from the enlarged overlay images for clarity. Scale bars display 20 μm or 5 μm. (D) Scatter plot showing the correlation between GFP-Gephyrin (from KI) and anti-Gephyrin staining intensity (from immunofluorescence). Correlation (slope) between GFP and anti-Gephyrin fluorescent intensity was calculated by Simple Linear Regression (GraphPad Prism 7 or 8). Data from 20 neurons and 1007 GFP-Gephyrin puncta.

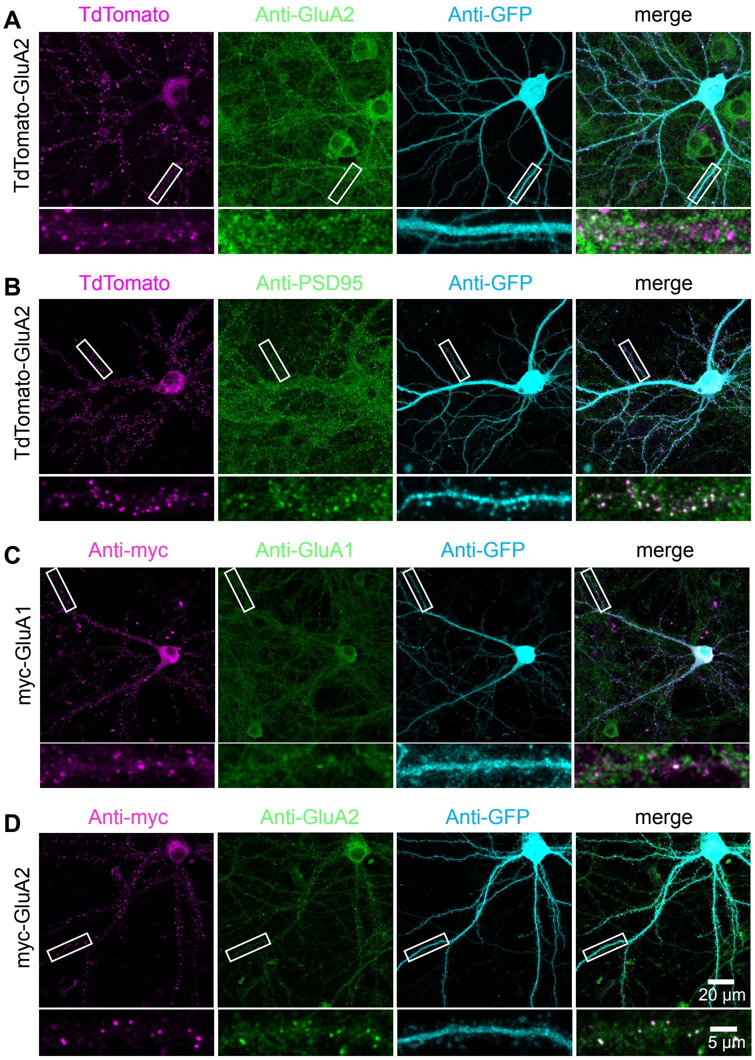

**Figure supplement 6. TKIT of alternative tags for AMPA receptors**

(A and B) Representative immunofluorescent images demonstrating TKIT of tdTomato-GluA2. The dendritic region within the white box is enlarged below to better illustrate co-localization between tdTomato-GluA2 (magenta) and GluA2 (green, A) or PSD95 (green, B). (C and D) Representative immunofluorescent images demonstrating TKIT of myc-GluA1 and myc-GluA2. The dendritic region within the white box is enlarged below to better illustrate co-localization between myc-GluA1 and myc-GluA2 (magenta) with GluA1 and GluA2 (green), respectively. Note: the cell-fill channel was removed from the enlarged merged images for clarity. Scale bars display 20 μm or 5 μm.

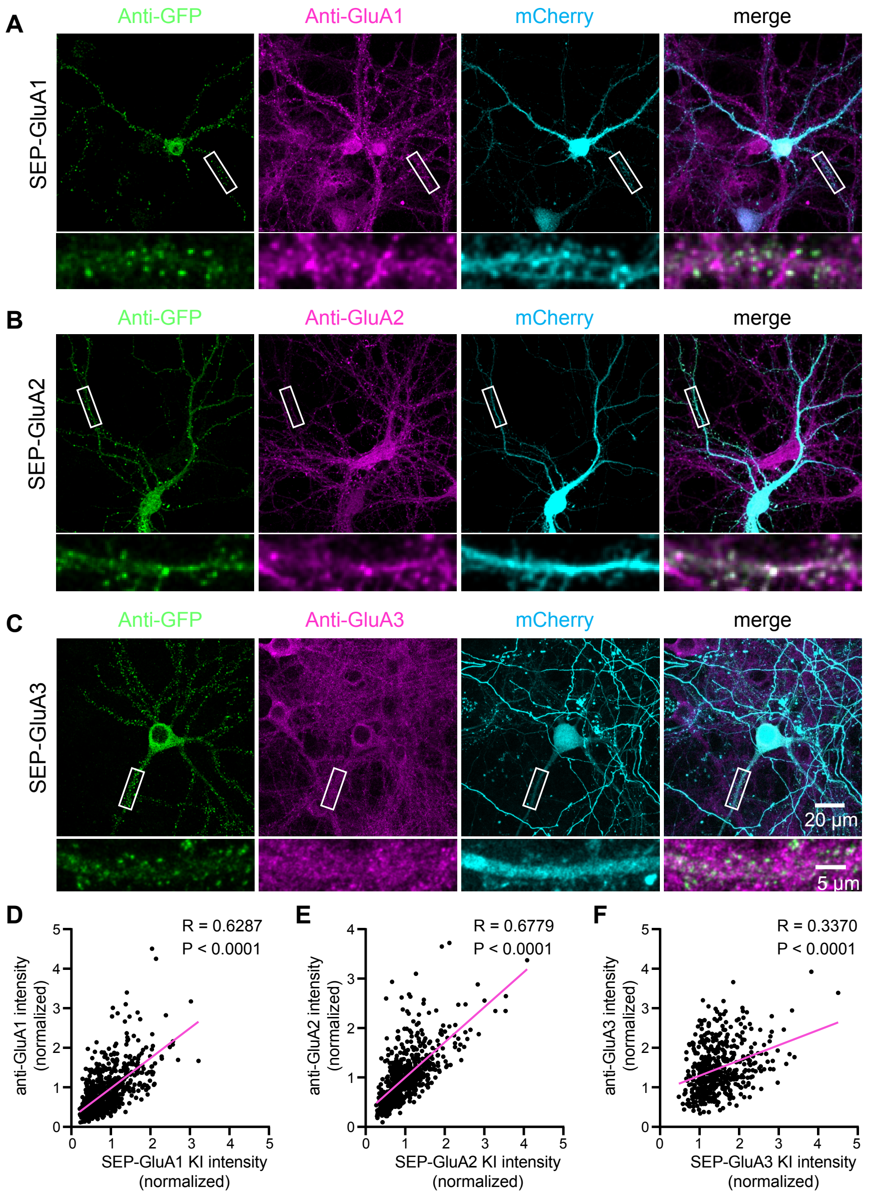

**Figure supplement 7. Correlation of SEP-GluA1-3 KI with endogenous AMPA receptors**

(A-C) Representative immunofluorescent images showing TKIT of SEP-GluA1 (A), SEP-GluA2 (B), and SEP-GluA3 (C). The dendritic region within the white box is enlarged below to better illustrate co-localization between SEP-GluA1 (green) and GluA1 (magenta, A), SEP-GluA2 (green) and GluA2 (magenta, B), and SEP-GluA3 (green) and GluA3 (magenta, C). Note: the cell-fill channel was removed from the enlarged overlay images for clarity. Scale bars display 20 µm or 5 µm. (D-F) Scatter plots showing the correlation between SEP-GluA1-3 and anti-GluA1-3 immunofluorescence intensity. Correlation (slope) between SEP and anti-GluAs fluorescent intensity was calculated by Simple Linear Regression (GraphPad Prism 7 or 8). Data from 14 neurons and 733 synapses for SEP-GluA1, 13 neurons and 668 synapses for SEP-GluA2, and 10 neurons and 508 synapses for SEP-GluA3.

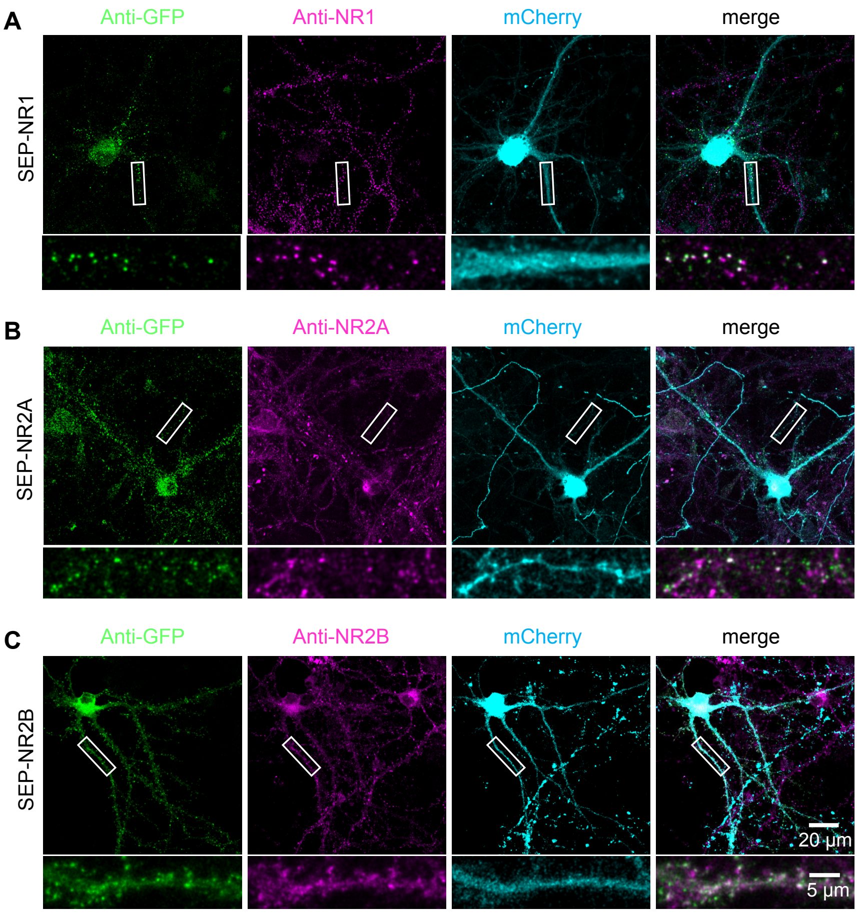

**Figure supplement 8. TKIT of SEP-NMDA receptors**

Representative immunofluorescent images demonstrating TKIT of SEP-NR1 (A), SEP-NR2A (B), and SEP-NR2B (C). The dendritic region within the white box is enlarged below to better illustrate co-localization between SEP-NR1 (green) and anti-NR1 (magenta, A), SEP-NR2A (green) and anti-NR2A (magenta, B), and SEP-NR2B (green) and anti-NR2B (magenta, C). Note: the cell-fill channel was removed from the enlarged overlay images for clarity. Scale bars display 20 μm or 5 μm.

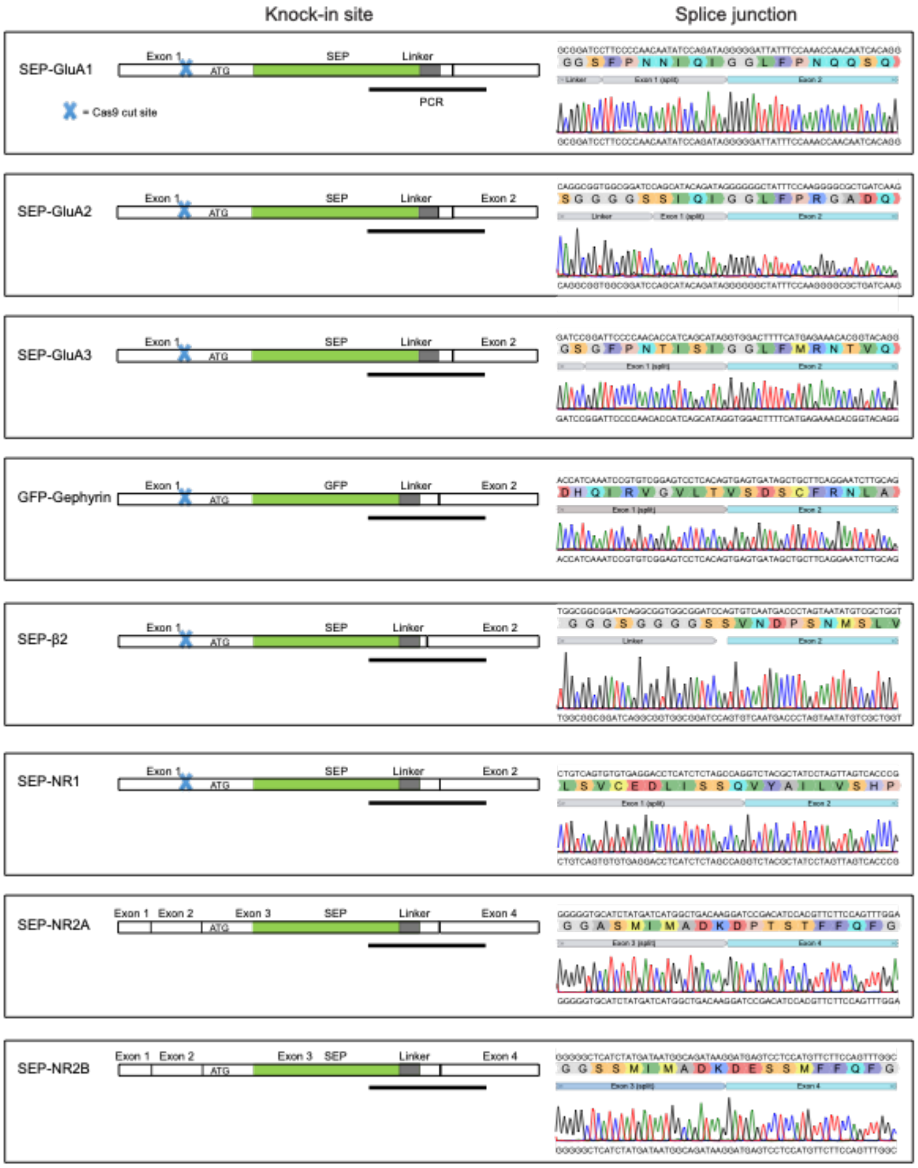

**Figure supplement 9. Intact splicing with TKIT**
Schematic of cDNAs indicating the KI site and neighboring exons for each SEP/GFP TKIT target. The blue cross indicates DNA cut sites. Sanger sequencing was used to analyze cDNA reverse-transcribed from mRNA extracted from neuronal cultures electroporated with each SEP/GFP TKIT target. Sequencing results indicate intact splicing of tagged proteins. The black line under the schematic indicate the region PCR amplified and sequenced. Note: the downstream cut site for SEP-GluA1, SEP-GluA2, SEP-GluA3, GFP-Gephyrin, SEP-β2, SEP-NR1 is located within introns and therefore is not indicated within the schematic. Both cut sites for SEP-NR2A and SEP-NR2B are located within introns.

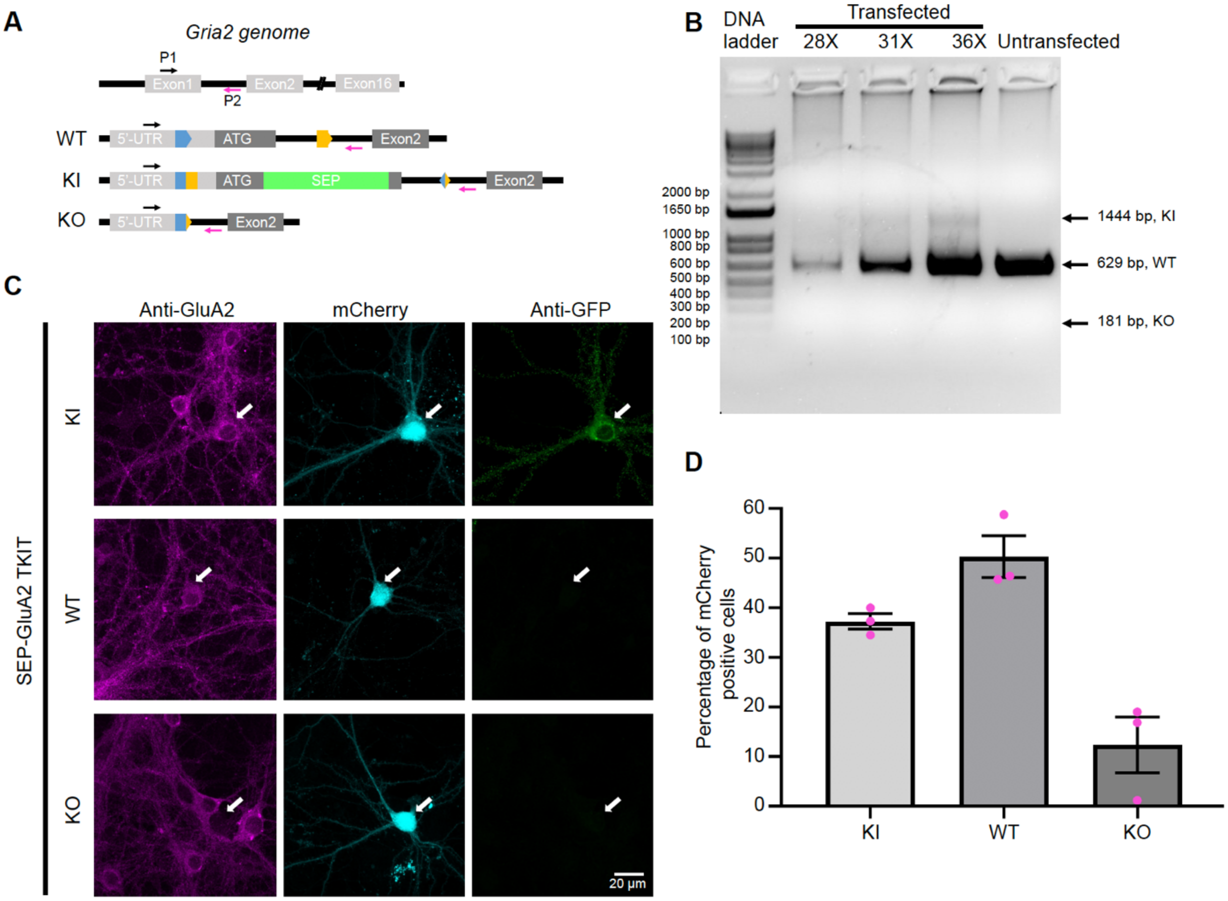

**Figure supplement 10. Low levels of GluA2 KO with TKIT**

(A) Top panel: cartoon showing *Gria2* gene structure and the location of PCR primers (upstream P1, black arrow and downstream P2, magenta arrow). Bottom panel: three predicted genotypes (WT, KI, KO). Arrows indicate the same primers showed in top panel. (B) Genomic DNA was extracted from neuronal cultures following electroporation of TKIT constructs. PCR was performed with one pair of PCR primers to detect KI/WT/KO outcomes with different numbers of cycles (28, 31, 36). Following DNA electrophoresis of PCR products, there was clear KI and WT bands, and no detectable KO band. Note: proportion of WT was overestimated, because part of WT was from non-transfected cells. (C) Representative immunofluorescent images of TKIT mediated SEP-GluA2 KI, including KI, WT and KO neurons. White arrows indicate transfected neurons. Scale bars represent 20 μm. (D) Quantification of different fractions of KI, WT, or KO cells based on data from 252 neurons (3 experiments) following SEP-GluA2 TKIT mediated KI. Mean of KI: 37.3%, WT: 50.3%, KO: 12.4%. Results shown as means ± SEM.

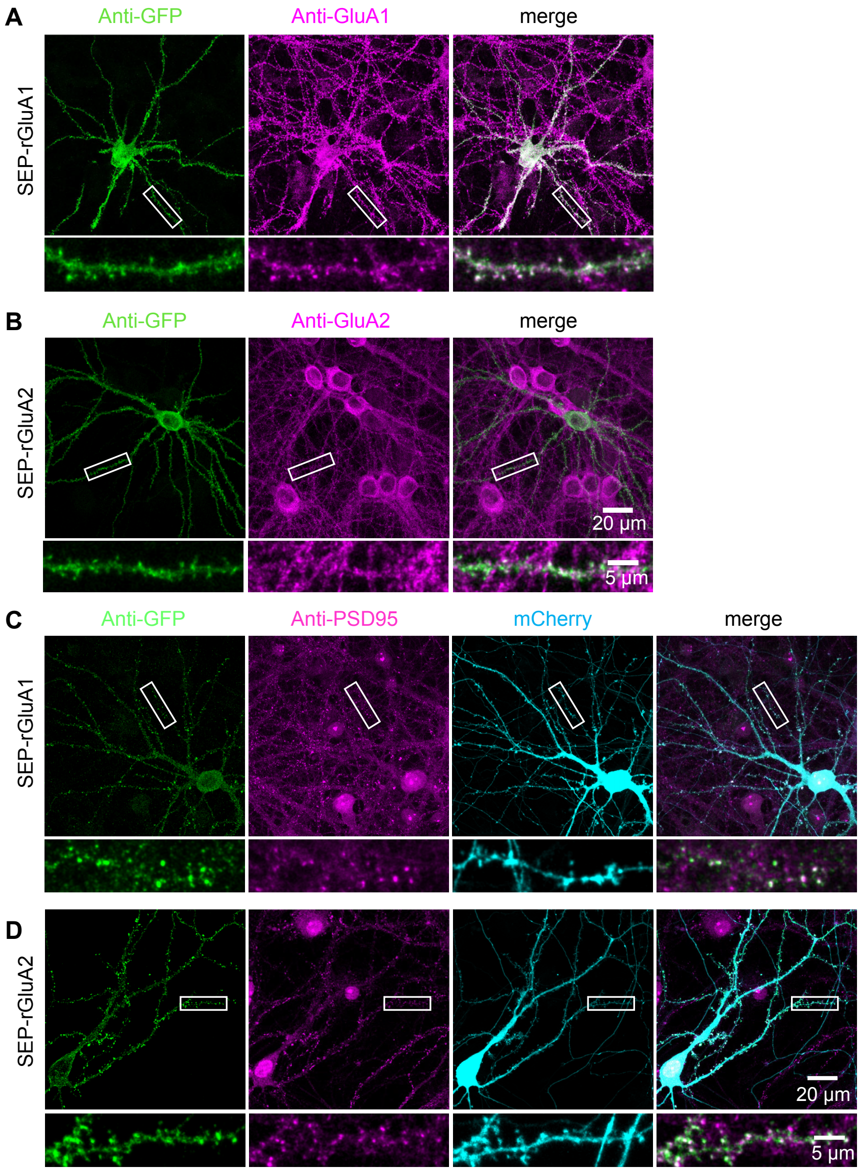

**Figure supplement 11. TKIT of SEP-GluA1 and SEP-GluA2 KI in rat neurons**

Representative immunofluorescent images demonstrating TKIT of SEP-GluA1 (A) and SEP-GluA2 (B) in rat primary cultured neurons. The dendritic region within the white box is enlarged below to better illustrate co-localization between (A) SEP-GluA1 (green) and GluA1 (magenta), or (B) SEP-GluA2 (green) and GluA2 (magenta). (C and D) Co-localization of SEP-GluA1 and SEP-GluA2 with PSD95. The dendritic region within the white box is enlarged below to better illustrate co-localization between (C) SEP-GluA1 (green) and PSD95 (magenta), or (D) SEP-GluA2 (green) with PSD95 (magenta). Note: the cell-fill channel was removed from the enlarged overlay images for clarity. Scale bars display 20 µm or 5 µm.

### Table supplement

| Gene | Guide1 | PAM | Guide2 | PAM | Guide 1 location | Guide 2 location |
|---|---|---|---|---|---|---|
| **MOUSE** | | | | | | |
| Gria1 | GGGAAGACCAAATCTATGGT | TGG | TTAGCAATGGAACACCAGGA | AGG | 5'-utr (exon1) | intron 1-2 |
| Gria2 | AACAGCCACCAGCTAAACCT | GGG | TATGCCTTTTGACACAATAG | AGG | 5'-utr (exon1) | intron 1-2 |
| Gria3 | GCGAGCGAGAGCAAGTTGAG | GGG | CTGCAAGAGGCTAAGAGTCG | GGG | 5'-utr (exon1) | intron 1-2 |
| Grin1 | GCGCTGCTCGAACACCCGCG | CGG | TCTGTTCCATCTAGTCAGTG | TGG | 5'-utr (exon1) | intron 1-2 |
| Grin2a | GCTCGGCTTGGACTGATACG | TGG | ACACCGAGCAATTCTCACAA | AGG | intron 2-3 | intron 3-4 |
| Grin2b | AAGAACCTCGTCTCCGTCTG | CGG | CATTGAATGACCCTTTCACA | GGG | intron 2-3 | intron 3-4 |
| Gphn | CGACCCCGAGTCCGGCCGGG | AGG | GAGAAACCTCCAGCAAGTCG | CGG | 5'-utr (exon1) | intron 1-2 |
| Gabrb2 | GGCGTACCAAAACATCAAAG | GGG | AGTCTGGTCACACTCTAAGG | GGG | 5'-utr (exon2) | intron 3-4 |
| **RAT** | | | | | | |
| Gria1 | GGGAAGACCAAATCTATGGT | TGG | CATGCTCGTCAAATGCCACG | AGG | 5'-utr (exon1) | intron 1-2 |
| Gria2 | ATATCGACCTCACAATGCAG | AGG | TATGCCTTTTGACACAATAG | CGG | 5'-utr (exon1) | intron 1-2 |

**Table supplement 1. Location and sequences of guides**

| | |
|---|---|
| **SEP-mGluA1** | |
| A1_fwd | GGCCTTCCTGGTGTTCCATTGCTAAGGTTGGACCAGGGCTTCT |
| A1_rev | GTGGGAAGACCAAATCTATGGTTGGTGGTGTTCCATTGCTAAGC |
| **SEP-mGluA2** | |
| A2_fwd | GGCCTCTATTGTGTCAAAAGGCATACCTGGGAGATAAGGATTC |
| A2_rev | GGAACAGCCACCAGCTAAACCTGGGTTGTGTCAAAAGGCATAGT |
| **SEP-mGluA3** | |
| A3.up_fwd | tccgcgcacatttccccgactcgagaattcccccatactcgtcagttctttgcAGGGGGTGTAAGAGCCGG |
| A3.up_rev | agccaccgccaccGGGGAATCCTCCGTGAGAATG |
| A3.SEP_fwd | cggaggattccccGGTGGCGGTGGCTCTAGAAG |
| A3.SEP_rev | tgttggggaatccGGATCCGCCACCGCCTGA |
| A3.dn_fwd | cggtggcggatccGGATTCCCCAACACCATC |
| A3.dn_rev | ctcgctgtatcgctcgagggatccgaattcgcgagcgagagcaagttgaggggAGCTCAGGTCTCCAATATC |
| **SEP-mNR1** | |
| N1.up_fwd | tccgcgcacatttccccgactcgagaattcccacactgactagatggaacagaCCAATACGCTTCAGCACCTCGGACAG |
| N1.up_rev | agccaccgccaccGGCAGCGCGGGCGAAGGA |
| N1.SEP_fwd | cgcccgcgctgccGGTGGCGGTGGCTCTAGAAG |
| N1.SEP_rev | cgcaggcagcgcgGGATCCGCCACCGCCTGA |
| N1.dn_fwd | cggtggcggatccCGCGCTGCCTGCGACCCC |
| N1.dn_rev | ctcgctgtatcgctcgagggatccgaattcccgcgcgggtgttcgagcagcgcTCACCAATGATAGTCACATGACAGCACACATGG TCC |
| **SEP-mNR2A** | |
| 2A.up_fwd | tccgcgcacatttccccgactcgagaattcacaccgagcaattctcacaaaggAGATGGACATGGGTCGAAGAATGTGG |
| 2A.up_rev | agccaccgccaccCTTCTCCGCCGCCGCGTT |
| 2A.SEP_fwd | ggcggcggagaagGGTGGCGGTGGCTCTAGAAG |
| 2A.SEP_rev | gagtacccttctcGGATCCGCCACCGCCTGA |
| 2A.dn_fwd | cggtggcggatccGAGAAGGGTACTCCAGCG |
| 2A.dn_rev | ctcgctgtatcgctcgagggatccgaattcgctcggcttggactgatacgtggTCAGAGGCCACTTAACCTG |
| **SEP-mNR2B** | |
| 2B.up_fwd | tccgcgcacatttccccgactcgagaattcccctgtgaaagggtcattcaatgGATTTTCTAGCATTCAGACACTTC |
| 2B.up_rev | agccaccgccaccGCTCTTTTGGGAACGAGC |
| 2B.SEP_fwd | ttcccaaaagagcGGTGGCGGTGGCTCTAGAAG |
| 2B.SEP_rev | gggcgctcttttgGGATCCGCCACCGCCTGA |
| 2B.dn_fwd | cggtggcggatccCAAAAGAGCGCCCCCAGC |
| 2B.dn_rev | ctcgctgtatcgctcgagggatccgaattcccgcagacggagacgaggttcttCCTGAATGGGTCACGACC |
| **SEP-mbeta2** | |
| Gb2.up_fwd | tccgcgcacatttccccgactcgagaattcagtctggtcacactctaaggggGTGCGCATGCGCTCCACA |
| Gb2.up_rev | agccaccgccaccATTGACACTAAAGAAAAAAATGACAATAACCAGGAATGAATAAAGAAG |
| Gb2.SEP_fwd | ctttagtgtcaatGGTGGCGGTGGCTCTAGAAG |
| Gb2.SEP_rev | ggtcattgacactGGATCCGCCACCGCCTGA |

| | |
|---|---|
| Gb2.dn_fwd | cggtggcggatccAGTGTCAATGACCCTAGTAATATG |
| Gb2.dn_rev | ctcgctgtatcgctcgagggatccgaattccccctttgatgttttggtacgccTCTTCTTGAAAGACCTCTC |
| **GFP-Gephyrin** | |
| gep.up_fwd | ctcgctgtatcgctcgagggatccgaattcgagaaacctccagcaagtcgcggCGCGGCCCGACTCCGCCC |
| gep.up_rev | cccttgctcactctagaCATGTTTCCCAGCGCAGTCACCGCAC |
| GFP_fwd | gctgggaaacatgtctagaGTGAGCAAGGGCGAGGAG |
| GFP_rev2 | cgagccaccgccaagCTTGTACAGCTCGTCCATGC |
| gep.dn_fwd | cgagctgtacaagcttggcggtggctcgggcggaggtgggtcaGCGACCGAGGGAATGATC |
| gep.dn_rev | tccgcgcacatttccccgactcgagaattccctcccggccggactcggggtcgCGGATAGTAGCCGTTGCG |
| **SEP-rGluA1** | |
| rA1.upSEP_fw d | gcacatttccccgactcgagcctcgtggcatttgacgagcatgGGTTGGACCAGGGCTTCTTTTTCG |
| rA1.upSEP_re v | ggaaattggcGGATCCGCCACCGCCTGA |
| rA1.dn_fwd | tggcggatccGCCAATTTCCCCAACAATATC |
| rA1.dn_rev | aggaagcggaagagcgcccagggaagaccaaatctatggttggGGCATTTGACGAGCATGAAAC |
| **SEP-rGluA2** | |
| rA2.up_fwd | tccgcgcacatttccccgactcgagccgctattgtgtcaaaaggcataCAGAGGATCTAATTTGCTG |
| rA2.up_rev | cgccaccactagaTATGCTGTTAGAAGAGACAC |
| rA2.SEP_fwd | ttctaacagcataTCTAGTGGTGGCGGTGGC |
| rA2.SEP_rev | tctgtatgctgttGGATCCGCCACCGCCTGA |
| rA2.dn_fwd | cggtggcggatccAACAGCATACAGATAGGTAG |
| rA2.dn_rev | gagcgaggaagcggaagagcgcccaatatcgacctcacaatgcagaggTTGTGTCAAAAGGCATAC |
| **PCR primers for AAV constructs** | |
| **SEP-GluA1 & SEP-GluA2** | |
| donor_fwd | ccatcactaggggttcctgcggccgcACATTTCCCCGACTCGAG |
| donor_rev | agcaaaaggccagCGCTCGAGGGATCCGAATTC |
| 2guides_fwd | ccctcgagcgCTGGCCTTTTGCTCACATG |
| 2guides_rev | ccatcactaggggttcctgcggccgcTTATGTAACGGGTACCACC |

**Table supplement 2. PCR primers for TKIT donors**
